# An unusual aspartic acid cluster in the reovirus attachment fiber σ1 mediates stability at low pH

**DOI:** 10.1101/2021.02.01.429088

**Authors:** Giulia Glorani, Max Ruwolt, Nicole Holton, Ursula Neu

## Abstract

The reovirus attachment protein σ1 mediates cell attachment and receptor binding and is thought to undergo conformational changes during viral disassembly. σ1 is a trimeric filamentous protein with an α-helical coil-coiled Tail, a triple β-spiral Body, and a globular Head. The Head domain features an unusual and conserved aspartic acid cluster at the trimer interface, which forms the only significant intra-trimer interactions in the Head, and must be protonated to allow trimer formation.

Here we show that all domains of σ1 are remarkably thermostable across a wide range of pH, even at the low pH of the stomach. Interestingly, we determine the optimal pH for stability to be between pH 5-6, a value close to the pH of the endosome and of the jejunum. The σ1 Head is stable at acidic and neutral pH, but detrimerizes at basic pH. When Asp^345^ in the aspartic acid cluster is mutated to asparagine, the σ1 Head loses stability at low pH and is more prone to detrimerize. Overall, the presence of the Body stabilizes the σ1 Head.

Our results confirm a role of the aspartic acid cluster as a pH-dependent molecular switch, and highlight its role in enhancing σ1 stability at low pH.

## INTRODUCTION

Mammalian orthoreovirus (reovirus) infects humans during their childhood (1, 2), causing mild respiratory and enteric infection (3, 4). So far, three different reovirus serotypes with different tropisms and hemagglutination profiles have been discovered: T1L (Lang), T2J (Jones), and T3D (Dearing) (5–9). Reovirus enters epithelial cells via a two-step mechanism, first binding sialylated carbohydrates with low affinity, and then engaging receptor ssuch as Junctional Adhesion Molecule A (JAM-A)orNogo-66receptor (NogoR1)with high affinity (10–13) (14). After uptake, reovirus is internalized into the endosome where it is subjected to proteolytic cleavage of the outer capsid, leading to the production of infectious subvirion particles (ISVPs) (15, 16). In particular, cleavage of the outer capsid protein μ1 leads to exposure of hydrophobic sequences that mediate membrane penetration (17, 18).

Cellular uptake is mediated by the attachment protein σ1: a trimeric, filamentous protein protruding from the vertices of reovirus (19, 20).σ1is anchored to the viral capsid at its N-terminus, followed by an α-helical coiled-coil region named Tail (21) and a tightly entwined Body domain featuring triple β—spiral repeats (22). The C-terminal Head region is a trimer of separately folded globular Head domains with a tight β—barrel fold, each composed of two Greek-key motifs (22). Both T1L and T3D σ1 engage JAM-A with a conserved binding site at the base of their Head domains (11) (23). However, the carbohydrate receptor on host cells and its binding domains on σ1 vary among different serotypes. The T3D Body domain binds to different terminal α-linked sialic acids (13, 22, 24, 25), while T1L σ1 engages the sialylated branched GM2 glycan with its Head domain (26).

Upon transition from virions to ISVPs, the attachment protein σ1 undergoes conformational changes (20) (27, 28) from a contracted form to a more elongated conformation, maybe the one observed in the crystal structures (22, 26, 29). However, neither the nature nor the trigger for these conformational changes is entirely understood. Candidates are carbohydrate binding (30), pH, and proteolytic cleavage (25). The binding of 9BG5 monoclonal antibody to virions but not to ISVPs, has been suggested as the evidence for a conformational change on the virion (20) (27). In addition, the binding of sialic acid >leads to higher affinity of the virus for JAM-A (30).

An attractive candidate for a molecular switch is the highly conserved cluster of aspartic acid residues at the trimer interface at the bottom of the globular Head (31, 32). In T3D σ1, three copies of aspartic acids Asp^345^ and Asp^346^each are buried in a hydrophobic and solvent-inaccessible pocket (10, 33).The side chains of protonated Asp^345^ form hydrogen bonds with the deprotonated side chains of Asp^346^ on the clockwise neighbouring monomer within the trimer. Evidence for the unusual protonation of Asp^345^ at neutral pH comes from a variant, in which Asp^345^ was replaced with neutral asparagine that cannot be deprotonated. The D345N σ1 Head formed trimers at neutral pH, bound JAM-A with similar affinity to the wild-type, and exhibited a highly similar overall structure (10).The aspartic acid cluster is shielded from the solvent by two layers of aromatic amino acids directly below and above the aspartic acid residues, likely protecting the protonated aspartic acid cluster at neutral pH. Both the aspartic acids and the neighbouring residues are highly conserved among different prototype and field isolate reovirus strains, indicating an important role for reovirus infection. Formation of the trimer of σ1 Head domains from monomers likely needs to take place in a relatively low pH environment that enables at least partial protonation of Asp^345^ in the monomer. Therefore, it is likely that these residues could play a role in inducing σ1 conformational changes, during the infection and the cell entry process (10, 33).

Reovirus encounters low pH in the endosome during infection of host cells, but it is also exposed to drastically varying pH during transmission and initial infection of a host organism. Following ingestion of contaminated food or inhalation of aerosols, reovirus travels to the gastrointestinal tract, encountering the stomach and the small and large intestines (34, 35). These environments provide different pH ranges, oxygen levels, and bacterial loads (36–38). While the pH is constantly acidic (1.5 – 3.5) in the stomach, the transition to the small and large intestine leads to shift towards mildly acidic and neutral pH ranges (6.3 – 7.3) (36, 39). Reovirus needs to withstand the low gastric pH and might use low pH as a trigger for conformational changes. However, there is no direct evidence on how changes in environmental pH can affect σ1 conformation, stability, and trimeric organization. In this study, we conducted a biophysical investigation on the thermostability and the oligomerization state of recombinant σ1 domains over a broad range of pH, including the very low pH of the stomach. We expressed and purified constructs spanning all of T3D σ1. We then determined their melting temperatures via differential scanning fluorimetry, identifying the pH of optimum stability of T3D σ1 constructs to be pH 5, a value close to the pH of the endosome and of the jejunum. We found by analytical size exclusion chromatography that the σ1 Head detrimerizes at high pH, confirming protonation of the aspartic acid cluster. The effect of the D345N mutation at the trimer interface was found to particularly affect the stability of σ1 trimer at neutral and low pH, upon incubation at room temperature. Altogether, our findings suggest that the aspartic acid at the trimer interface of the σ1 Head domain is required for the stability of the trimeric Head domain, especially in a low pH environment.

## EXPERIMENTAL PROCEDURES

### Protein expression and purification

T3D σ1 Tail-Body (25-291) construct was a kind gift of the Stehle laboratory (UniversitätTäbingen). T3D σ1 Tail-Body was subcloned into pETM11 vector in frame with a N-terminal tag consisting of a hexahistidine (His) motif for affinity purification and a Tobacco Etch Virus (TEV) protease cleavage site for removal of the His-tag during purification (40). After cleavage of Tail-Body, the amino acids GAMA remained at the N-terminus.

Body-Head (170-455) and Head (293-455) constructs, which lack the stably trimeric Tail, had been subcloned into a modified pQE80L vector and additionally provided with a trimeric version of the GCN4 leucine zipper to induce trimerization (41, 42). After cleavage, the amino acids GAMA remained at the N-terminus of the cleaved protein. Site-directed mutagenesis was used to produce T3D D345N σ1 in the context of the Body-Head and Head constructs.

Upon transformation into BL21(DE3)-RIL cells, protein was expressed by autoinduction (43) at 20 °C for 4 days. Bacteria were harvested by centrifugation and the bacterial pellet was resuspended in 40 mM HEPES pH 7.1, 300 mM NaCl, and 10 mM Imidazole, supplied with 1 mM PMSF, 0.01 mg/mL DNase I, and 0.2 mg/mL Lysozyme. Cells were lysed with an Emulsiflex-C5 Homogenizer (Avestin) and the insoluble fraction was removed by centrifugation at 21,500 X g. The soluble fraction was purified via immobilized metal affinity chromatography (IMAC) on a HisTrap^TM^FF column (GE Healthcare) and eluted with a linear gradient to 40 mM HEPES pH 7.1, 300 mM NaCl, 500 mM Imidazole. Controlled TEV-protease treatment was performed during dialysis against 40 mM HEPES pH 7.1, 300 mM NaCl, 10 mM Imidazole at room temperature for 4 hours, in order to cleave the 6X-HisTag and simultaneously reduce the Imidazole concentration. The cleaved protein was isolated via reverse mode IMAC and further purified by gel filtration on a HiLoad® 16/600 Superdex® 200 pg in 40 mM HEPES pH 7.1, 300 mM NaCl. The pure protein was either used freshly or cryoprotected by the addition of 20% (v/v) ethylene glycol, flash-frozen in liquid nitrogen and stored at –80 °C.

### Differential scanning fluorimetry

Differential scanning fluorimetry was performed adapting the protocol from Niesen F.H. et al. (44). T3D σ1 constructs were thawed and dialyzed multiple times on ice with a ZelluTrans/Roth Mini Dialyzer (Roth) against 40 mM HEPES pH 7.1,150 mM NaCl, to remove the cryo-protectant. They were then diluted to 0.2 mg/mL and mixed with SYPRO orange (Sigma-Aldrich) to give a final dye:protein molar ratio of 1:500. T3D σ1 constructs were tested against buffers in a range of pH between 1.4 and 9.4 (Table S1). In a 96-well plate, 10 μL of the different assay buffers (0.4 M) were mixed with 10 μL of σ1 protein to give final buffer and protein concentrations of 0.2 M and 0.1 mg/mL, respectively. Melting curves were recorded in triplicate on a Mx 300P qPCR system (Agilent), between 25 °C and 95 °C, with a scanning rate of 1 °C/min. Data was fitted to the Boltzmann equation with Origin® Pro 9.5.

### Analytical gel filtration

T3D σ1 constructs were dialyzed at room temperature with ZelluTrans/Roth Mini Dialyzers (Roth) against buffers titrated to pH 3.0, 5.0, 7.1 and 8.6 (Table S2), to eliminate the cryo-protectant and equilibrate the sample to the assaypH. T3D σ1 samples were tested before and after an additional 4-hour incubation step at room temperature. 50 μL of 0.4 mg/mL σ1 protein were run on a Superdex® 200 Increase 3.2/300, at a flow rate of 0.075 mL/min while monitoring the protein absorbance at 280 nm. Elution peaks corresponding to trimers and monomers were integrated in Unicorn (GE Healthcare, Cytiva).

## RESULTS

### T3D σ1 is stable over a wide range of pH values

To determine the effect of pH on the stability of different σ1 domains, we recombinantly expressed and purified three different constructs spanning the entire length of σ1 (Fig. 1A): a Tail-Body construct spanning the coiled-coil Tail as well as most of the Body domain (25-291), a Body-Head construct containing the entire body domain as well as the Head domain (170-455) and the Head domain with the last β-spiral (293-455). All σ1 wild-type constructs were purified as trimers at pH 7.1. To assess the effect of pH on protein stability, we incubated the σ1 constructs with an excess of buffers of different pH ranging from 1.4 to 9.4 in increments of 0.5 pH units (Table S1) and acquired thermal denaturation curves at each pH by differential scanning fluorimetry. While most proteins evolved against thermal denaturation under physiological conditions, the melting temperature reports on the Overall stability of a protein domain – the higher the melting temperature, the more rigid and stable the protein domain. Thus, melting temperatures also report on changes in protein conformations or dynamics under different conditions. As reovirus travels through the stomach and gastrointestinal tract, we hypothesized that σ1 constructs might be more stable in a low pH environment.

**Figure 1:**
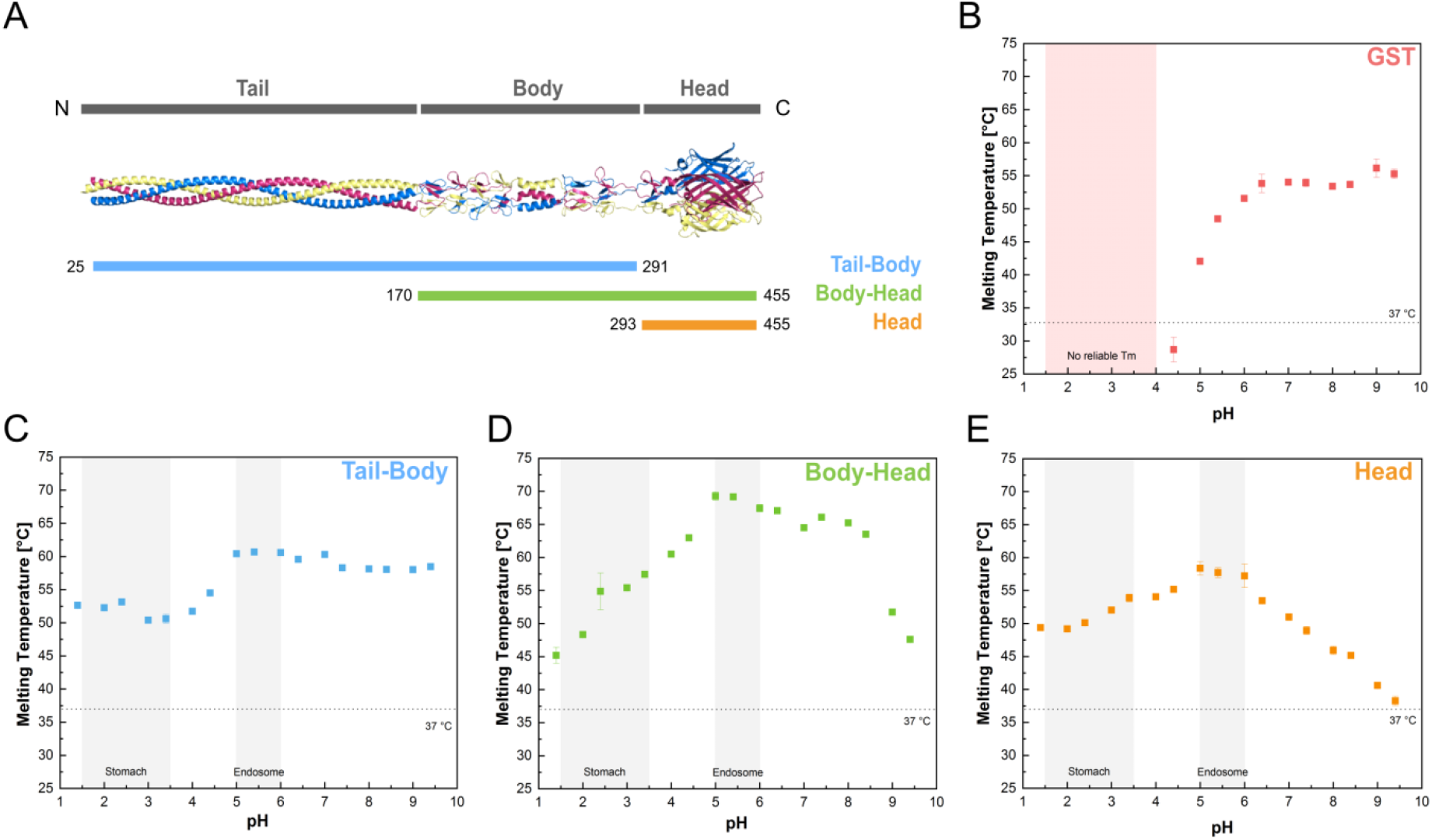
A) Structural model and domain organisation of reovirusσ1. The model was generated by superposition of overlapping published crystal structures (pdb codes 3s6x and 6gap)(21, 22). The trimeric protein is shown in cartoon representation and coloured according to chain. The constructs used in this study are colour coded according to their length (Tail-Body in light blue, Body-Head in green, and Head inorange). B-E) Thermostability of T3D σ1 Tail-Body (C), Body-Head (D) and Head (E) constructs in buffers of different pH, with GST as comparison (B). Melting temperatures are plotted as averages of three independent experiments, with error bars showing standard deviation. The range of pH of stomach and endosome are highlighted in light grey. For GST, no melting temperatures were obtained below pH 4.4 because the protein denatured upon mixing with the low pH buffers before starting the experiment.

All T3D σ1 constructs featured high thermostability at low pH values between 1.4 and 5, with melting temperatures between 45 and 70 °C (Fig. 1C-E). In particular, all constructs have a melting temperature much higher than 37 °C even at the highly acidic pH of pH 1.4, with 53, 45 and 50 °C for the Tail-Body, Body-Head, and Head constructs, respectively. For comparison, the acid-instable model protein Glutathion-S-transferase (GST) instantly denatured at temperatures lower than 25 °C at pH values of 4 and below (Fig. 1B). The unfolding signal of the Tail-Body construct is strongly determined by unfolding of the Body domain as the Tail domain does not contain aromatic amino acids to which the dye used for detection could bind.

When comparing melting temperatures across the pH spectrum, the Tail-Body construct featured a single transition, with constant melting temperatures of roughly 50 °C (between pH 1.4 and 4, and constant melting temperatures of roughly 60 °C between pH 5 and 9.4 (Fig. 1C). In contrast, the Head domain featured a stability optimum of 58 °C between pH 5 and 6, with a gentle drop in stability at lower pH and a steep drop in stability at !higher pH, leading to a melting temperature of only 38 °Cat pH 9.4 (Fig. 1E). A melting temperature of 38 °C was the lowest value acquired among all the constructs tested and the closest to the physiological temperature of the human body. The Body-Head construct was more stable than the Tail-Body or the Head constructs, with a broad pH optimum of stability of 67-70 °C between pH 5 and 8.4, while Tail-Body and Head constructs featured 10 °C ‘lower melting temperatures at pH 5. In addition, the Body-Head construct featured deep drops in stability towards both low and high pH: a moderate drop towards lower pH values and a steep drop to about 48 degrees between pH 8.5 and 9 (Fig. 1D). When tested for its thermostability in a various range of pH, the T3D σ1 Body-Head construct showed overall higher melting temperatures than the Head domain expressed alone. Although the melting temperature values were comparable between the two constructs in the low pH of the stomach, at the optimum pH of 5, their difference was about 10 °C. The same gap in melting temperatures could also be detected upon incubation with basic pH environment. In addition, from pH 5 onwards, the Head construct showed a sharp decreasing in melting temperatures, reaching its lowest value of 37 °C, at pH 9.4. Interestingly, when the Body domain was ‘expressed together with the Head domain, its overall stability was enhanced and the lowest melting temperature detected was 47 °C, at pH 9.4.

Taken together, the stability optimum of all constructs lays around pH 5-6, which (interestingly coincides with the pH of the endosome as well of the upper gut. In addition, while all T3D σ1 constructs were stable at acidic pH, there were differences between the constructs upon incubation in neutral and basic buffers. Consequently, we conclude that the different domains of the elongated σ1 molecule are differentially affected by changes in environmental pH. While the increase in pH did not influence the melting temperature of the Body-Tail construct, it caused significant destabilization of the Head and Body-Head constructs. We therefore hypothesized that the high-pH drop in thermostability of the constructs containing the Head domain was caused by the Head domain, which features an aspartic acid cluster at the trimer interface that had been proposed to act as a pH-sensing switch(10).

### The Head domain of T3D σ1 detrimerizes at high pH

The highly conserved aspartic acid cluster and its surrounding structures at the bottom of the σ1 Head domain are the main trimerization interface of the σ1Head(10). Based on the high-resolution crystal structure of the σ1 Head, it was hypothesized that the trimer of σ1 Head domain can only be formed and maintained if Asp^345^ in the cluster interface is protonated. At neutral pH, protonation of Asp^345^can only be maintained by shielding it from solvent. However, if thermal motion or a basic environmental pH overcome the shield and Asp^345^ is then deprotonated, repulsion between the negatively charged Asp^345^ residues would cause the σ1 Head to detrimerize.

Based on our thermostability data, which showed changes in stability with pH, we therefore investigated the multimeric state of the σ1 Head domain at different pH values by analytical size exclusion chromatography, which reports on the hydrodynamic radius of macromolecules. We used homogeneous purified trimeric σ1 Head and incubated it for 4 hours at RT in the respective pH before analysis. After analytical size exclusion chromatography, the amount of monomeric T3D σ1 was calculated as a fraction of the total loaded protein. At acidic and neutral pH values, the wild-type Head construct was almost exclusively trimeric, even after incubation for 4 hrs at RT. At pH 8.6, the wild-type Head partially dissociated into monomers, leading to a 25% fraction of monomeric wild-type Head. Our results without incubation are in accord with previous findings indicating the wild-type σ1 Head to be trimeric when analysed at 4°C and without incubation (10). However, our experiments were performed at 20 °C or higher, which more closely resembles physiological conditions and provides increased thermal motion leading to more dynamic protein structures (45).

### D345N T3D σ1 mutant Head is overall less thermostable than the wild-type Head

To investigate the role of the protonated aspartic acid cluster in T3D σ1 Head stability, we replaced Asp^345^ with an asparagine to mimic a permanently protonated aspartic acid.

While aspartic acid features a carboxylic acid group (-COOH), asparagine carries an amide !group (-CONH_2_) in which one of the oxygens is replaced with a nitrogen atom. Due to this chemical difference, free aspartic acid is deprotonated and negatively charged at neutral pH, while asparagine is not deprotonated even at basic pH. Moreover, asparagine can engage in the same pattern of hydrogen bonds as protonated aspartic acid.As asparagine cannot be deprotonated, we anticipated the D345N σ1Head to be more resistant to thermal denaturation and essentially morestable at the trimer interface than the wild-type at neutral to basic pH.

The D345N σ1 Head construct featured similar melting temperatures to that of the wild-type, with roughly 43°C at mildly acidic, neutral and basic pH. However, the D345N σ1 Head construct showed no drop-off in stability at basic pH, in contrast to the wild-type (Fig. 2A). The finding confirmed our hypothesis on the role of D345. However, unexpectedly, the overall melting temperatures of the D345N mutant Head were much lower than those of the wild-type construct across pH 1.4 to pH 8.4. Compared to the fairly stable wild-type Head, D345N Head had a 10-15 °C lower melting temperature at mildly acidic and neutral pH, and ia 25 °C lower melting temperature at very low pH. In particular at pH 1.4-3.4, the melting temperatures of the D345N mutant were 10°C below physiological temperature, indicating that it might be instable at gastric pH. Meanwhile, the wild-type Head construct was fairly thermostable at very low pH.

**Figure 2:**
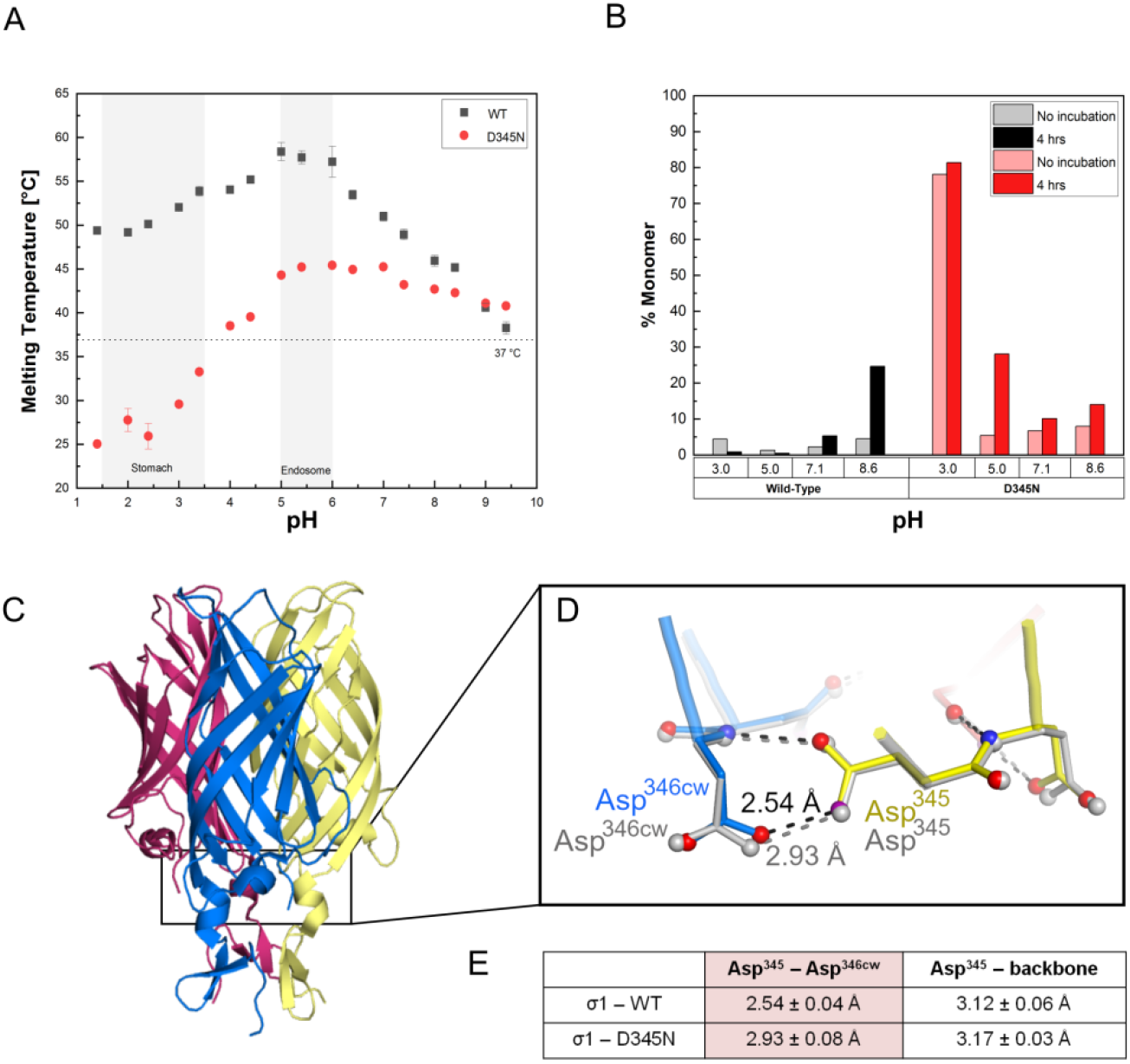
A) Thermostability of T3D σ1 wild-type (black) and D345N (red) mutant Head. The range of pH of stomach and endosome are highlighted in light grey. Data is an average of three independent experiments, error bars represent the associated standard deviation. B) Analytical size exclusion chromatography of wild-type (grey and black) and D345N mutant Head (light red and red). Percentage of monomeric T3D σ1 Head is depicted after integrating absorbance peaks from the chromatograms (Fig. S1). Measurements were carried out directly after buffer exchange (light colour) and after a 4-hr incubation at room temperature (darker colour). C) Location of the aspartic acid cluster in the σ1 Head. σ1 is shown in cartoon representation, with one monomer coloured yellow, one blue and one red. Figure is based on pdb entry2OJ5 (10). D) Organisation of the aspartic acid cluster in the T3D σ1. Thewild-type protein is shown in stick representation and coloured yellow for one chain and blue for its clockwise neighbouring monomer. The D345N mutant is coloured grey. The hydrogen bonds from Asp345 to Asp346cw are shown as dashed lines for the wild-type (black) and D345N (grey) structures. Figure is based on pdb entries 2OJ5 and 2OJ6 (10). Atomic distances between residues were measured with Coot. E) Atomic distances between side chain oxygen and nitrogen atoms of Asp345 (wild-type) or Asn345 (mutant) to their hydrogen bonding partners. Distances were averaged over the six crystallographically independent monomers in the structures and given with their standard deviations.

### D345N T3D σ1 Head detrimerizes into monomers

We then asked whether the decreased thermostability of the D345N Head correlated with its multimeric state and performed analytical gel filtration at different pH values. At mildly acidic and neutral pH, a 10 % fraction of the D345N mutant Head dissociated into monomers (Fig. 2A). Interestingly, at pH 3.0, D345N had completely dissociated into monomers during the buffer exchange and no further detrimerization was observed during incubation at room temperature. In contrast, the wild-type constructs had been almost exclusively trimeric at acidic and neutral pH (Fig. 2A). The decreased stability of the D345N Head trimer at acidic pH correlates with the decreased thermostability of the protein in the same pH range. Moreover, the wild-type protein was predominantly trimeric at acidic pH and much more thermostable. Therefore, the D345N mutation likely caused the decreased thermostability by disrupting the trimerization interface at acidic pH. Our results therefore point to a role for the highly conserved aspartic acid in the trimerization interface in stabilizing the trimer at low pH.

### Structural basis for the lower stability of D345N mutant Head

The D345N mutation did not change the charge, the packing of side chains, nor the pattern of hydrogen bonds as D345 is protonated, but it did alter charges in the trimer [interface. Yet, the variation had a strong effect on trimer integrity and overall protein stability. To elucidate its mechanism, we compared the previously determined high-resolution crystal structures of wild-type and D345N σ1Head (PDB accession codes: 2OJ5 and 2OJ6, respectively)(10).

The wild-type and D345N proteins had crystallized in the same space group with identical unit cell parameters, the same two trimers in the asymmetric unit and diffracted to similar resolution of 1.75 and 1.85 Å resolution, respectively. The overall structures were virtually identical with main chain r.m.s.d. value of 0.13 Å, consistent with the fact that the two proteins differ in only one heteroatom. However, there was a significant difference between the two variants in the distance between the side chain of residue 345 and its hydrogen (bonding partner across the trimer interface, Asp^346cw^ (cw indicating the clockwise neighboring (monomer in the σ1 trimer). This distance averaged 2.54 Å among the six crystallographically (independent monomers in the asymmetric unit in the wild-type crystal, but it averaged 2.93Å in the D345N crystal (Fig. 2C). Thus, this hydrogen bond was 0.4 Å longer in the D345N (mutant than in the wild-type protein (statistically significant with p=1.7*10^-5^ by Welch t-test). (The length of hydrogen bonds inversely correlates with their strength (46–50), with 2.5 Å indicating a very strong bond and 3.5 Å often used as a generous cutoff for weak hydrogen >bonds. Thus, the wild-type trimer is held together by a much stronger hydrogen bond than the D345N mutant.

### The addition of the Body to the Head domain enhances stability

On the virus, the globular σ1Head domain is linked to the σ1 Body domain, which overall stabilized the Head domain (Fig. 1D,E) and might counteract the D345N mutation. We therefore tested the effect of the D345N mutation on the stability of the Body-Head construct. Similar to the D345N Head alone, the overall thermostability of D345N Body-Head did not decrease at high pH (Fig. 3A), while the thermostability of the wild-type Body-Head construct was reduced in a basic environment. However, the overall melting temperatures of the D345N mutant Body-Head were 10 °C lower than those of the wild-type construct at acidic and neutral pH, confirming the stabilizing effect of Asp^345^ in the wild-type protein. Again, the D345N mutant Body-Head construct was least stable at very low pH, reaching its lowest value of 32 °C at pH 1.4. When comparing D345N Head and Body-Head domains, the addition of the Body to the Head domain stabilized the protein overall to give 15 °C higher melting temperatures across the pH range tested.

**Figure 3:**
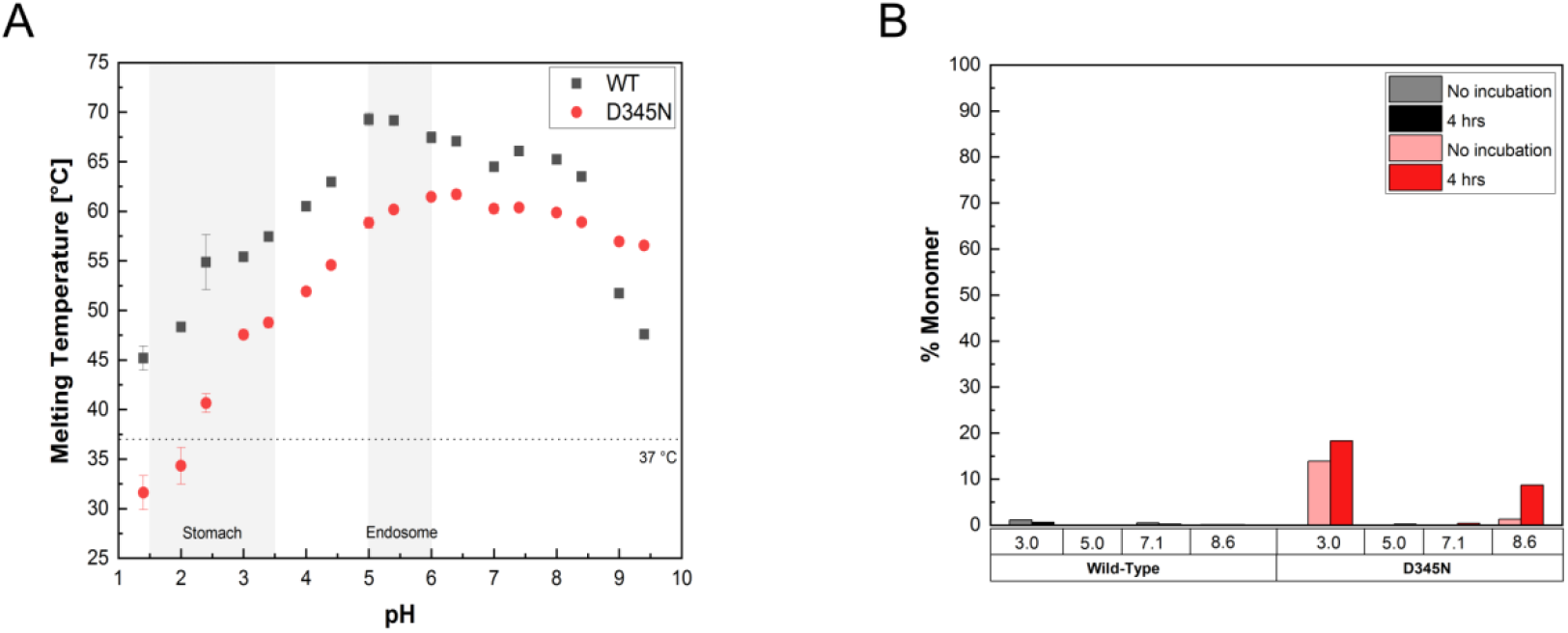
A) Thermostability of T3D σ1 wild-type (black) and D345N (red) mutant Body-Head construct. The range of pH of stomach and endosome are highlighted in light grey. Data is an average ofthree independent experiments, error bars represent the associated standard deviation. B) Analytical size exclusion chromatography of wild-type (grey and black) and D345N mutant Body-Head (light red and red). Percentage of monomeric T3D σ1 Body-Head is indicated after integrating absorbance peaks from the chromatogram. Measurements were carried out before (light color) and after a 4-hour incubation time (dark color).

We then asked whether the addition of the Body domain prevented detrimerization of the wild-type and D345N Head at low pH and determined its oligomeric state by analytical size exclusion chromatography. Interestingly, the wild-type Body-Head construct was present almost exclusively as a trimer after incubation in different pH buffers, both directly after buffer exchange and after 4 hrs incubation at room temperature (Fig. 3B). The D345N Body-Head construct showed some detrimerization at pH 3.0 (15% monomer) and after 4 hrs incubation at pH 8.6 (8% monomer). Therefore, the Body domain prevented detrimerization of the wild-type protein at high pH (Fig. 2B, 3B) and greatly reduced detrimerization of D345N at low pH.

## DISCUSSION

σ1 is thought to adopt a contracted conformation on the virion and is hypothesized to undergo a conformational change to a more elongated form during virus entry (20, 27). An antibody against the outer capsid protein σ3 prevents hemagglutination of red blood cells, which is mediated by the sialic acid binding site located in the body domain of σ1. Thus, either the Body or the Head domain of σ1 is likely close to the virus surface in the contracted form. The recombinant σ1 domains used in this study are all elongated (22, 29) and therefore are thought to represent the structure of σ1 after the conformational change. While the Body domain seems to feature a relatively flat pH profile of thermostability, the σ1 Head is the only domain whose thermostabilityis strongly influenced by pH. Even at strongly acidic pH, wild-type σ1 Head is a highly stable trimer, with no detrimerization occurring and a pH optimum of thermostability of the protein between pH 5 and pH 6. This is likely an adaptation of the protein structure to the exposure to low intragastric pH during transmission. Similarly, structural studies of the enteric adenovirus Ad-F41 short fiber head revealed a loop in the fiber head that was disordered at neutral pH and ordered at pH 5 (51, 52). Moreover, comparison of the Ad-F41 capsid to that of non-enteric adenoviruses highlighted a surface with relatively few charged residues and an almost unaltered structure at pH 4 (53)

We find here that the thermal motion at room temperature combined with a basic pH of 8.6 causes slow detrimerization of the wild-type σ1 Head. This result directly confirms in solution earlier conclusions from the crystal (10, 33) that the σ1 Head trimer might be labile to high pH. Earlier molecular dynamics simulations suggested that deprotonation of Asp^345^ causes opening of the Head trimer by electrostatic repulsion (33). Moreover, removing the hydrophobic shielding from the aspartic acid cluster of σ1 led to purification of a monomeric σ1 Head construct, likely by allowing direct deprotonation of Asp^345^(10). Thus, a likely mechanism for detrimerization could be that the σ1 Head undergoes opening and closing “breathing” motions due to thermal motion, which might lead to deprotonation of Asp^345^ at basic pH. This model is consistent with the crystal structure featuring solid hydrophobic interactions between monomers at the bottom of the Head and weaker interactions at the top of the Head (10). However, at acidic and neutral pH, complete detrimerization of the wild-type σ1 Head was negligible, indicating that the trimer was stable under those conditions. This result, however, does not rule out that the σ1 Head might undergo opening and closing breathing motions at neutral without deprotonation of Asp^345^ and complete detrimerization. Our gel filtration assay cannot distinguish between open and closed trimer conformations of a small domain such as the σ1 Head.

To investigate the role of Asp^345^ as a pH-dependent switch, we then replaced Asp^345^ at the centre of the trimer interface with asparagine, which cannot be deprotonated and engages in the same pattern of hydrogen bonds as protonated aspartic acid. The biggest effects of the mutation were observed at acidic pH: a considerable decrease in overall thermostability correlating with an increased tendency to detrimerize. Even at its pH optimum of stability, the D345N mutant had a 15 °C lower melting temperature than the wild-type. This effect was even more pronounced at pH 3, which the virus might encounter in the stomach.

As the stability of the D345N trimer depended on the pH of incubation, it is likely that there are ionizable residues in the σ1 Head that destabilize the trimer at lowpH. One candidate is His349, three copies of which are located on the inner surface of the trimer just above the top of the aspartic acid sandwich and whose pK value lies in the relevant pH range. It is thus likely that the strong interactions in the aspartic acid sandwich hold the trimer together against pH-dependent repulsion in other parts of the protein. This hypothesis argues for an equilibrium of forces at the trimer interface of σ1.

Inspection of the previously solved wild-type and D345N crystal structures provide a framework to understand the observed differences in trimer stability (10). The -COOH to ^-^ OOC-hydrogen bond mediated by the protonated Asp^345^ to the deprotonated side chain of Asp^346^ on the clockwise neighbouring monomer within the trimer is quite short, with a distance between the two oxygen atoms involved of 2.5 Å. Short hydrogen bonds correlate with increased bonding strength (54). In the D345N mutant, the corresponding -CONH_2_ to ^-^ OOC-hydrogen bond is 0.4Å longer, which correlates with a much weaker hydrogen bond (54). Thus, it is likely that a protonated D345 can hold a marginally stable trimer together, even at low pH and elevated temperatures, while the bonding strength of N345 might not be enough. Interestingly, the corresponding bond in the 2.2 Å crystal structure of the T1L σ1 Head (55) is even shorter with an average length of 2.4 Å. Thus, the increased bonding strength of the aspartic acid cluster is likely a conserved feature among different reovirus serotypes.

When the Head domain was expressed together with the Body portion, we observed enhanced thermostability across the pH range. Likewise, the Body-Head domain seemed to almost fully prevent detrimerization of the Head domain. We envisage a model in which a relatively pH-invariant Body domain prevents detrimerization of the Head, which is more sensitive to changes in pH. While the Head purified alone can detrimerize readily according to pH, Body-Head trimers might undergo motions of the Head domain, but do not detrimerize completely when linked to the Body domain.

We find that lower thermostability of σ1 head correlated with a higher propensity to detrimerize into monomers. This result suggests that monomeric σ1 Head is less thermostable than the trimer. Interestingly, a σ1 Head mutant that was purified as a monomer was unable to recognize JAM-A (10) even though JAM-A contacts only one σ1 monomer and the mutation was not in the σ1-JAM-A interface. These results can be explained with decreased stability of the monomeric mutant causing a loss of function. Likewise, in the context of the virus, T1L Reovirus bearing disulfide crosslinks between the monomers in the Head domain had higher avidity for its receptor JAM-A than virus without crosslinks (56). Furthermore, enhanced stability of the σ1 trimer at the Head domain correlated with increased plaque size. Presumably, σ1 on virus without crosslinks can partially detrimerize at the Head, with a loss of stability and reduction in JAM-A binding. Moreover, not all crosslinked viruses featured all three possible crosslinks in the Head domain, indicating that the Head domain was not as rigidly trimeric as the body or tail domains.

Trimer-monomer dynamics have lately been observed for several trimeric viral surface proteins, most prominently pre-fusion conformations of viral class I fusion proteins. For example, antibodies which target the inner surfaces of the trimer interface of influenzavirus hemagglutinin are produced in infected and vaccinated individuals and confer protection in vivo. Binding assays suggest that the inter-trimer interface is at least temporarily accessible to these antibodies and can be disrupted by them, most prominently in the uncleaved HA0 trimer (57–59). Moreover, the SARS-CoV-2 Spike protein can exist in closed and open conformations, depending on whether the receptor-binding domains are buried in the protein or facing up towards the cell membrane. Interestingly, these dynamics are influenced by pH, with one or more receptor-binding domains facing up at neutral pH, while all of them facing down towards the protein at pH 4.5 (60), preventing binding of neutralizing antibodies against the receptor-binding domain. The protonation of several aspartate residues in a hinge region seems to play a role in this conformational change, but no cluster of aspartate residues is present in the SARS-CoV-2 spike.

Taken together, our results point to a role for the reovirus aspartic acid cluster not only as a potential pH-dependent molecular switch, but also as a requirement for stability of the σ1 Head at the low gastric pH all enteric viruses are exposed to and for the integrity of the JAM-A binding site.

## ACKNOWLEDGMENTS

We thank members of the Neu lab for helpful discussions and Svearike Oeverdieck and Daria Ivashinenko for help with protein purification. We thank Thilo Stehle (University of Tübingen) for his gift of initial σ1 expression plasmids and for fruitful discussions. We thank Markus Wahl (Freie Universität Berlin) for providing the research environment and for his insight into strong hydrogen bonds. This research was funded by the Emmy Noether program of the German Research Foundation (grant no. NE2017-1/1).

---

**Supplemental Table 1:** Buffers used during DSF measurements.

| pH | Buffer |
| --- | --- |
| 1.4 | 0.4 M HCl/KCl, 150 mM NaCl |
| 2.0, 2.4, 3.0 | 0.4 M Glycine/HCl, 150 mM NaCl |
| 3.4 | 0.4 M Citric Acid/Sodium Citrate, 150 mMNaCl |
| 4.0, 4.4 | 0.4 M NaAcetate, 150 mMNaCl |
| 5.0, 5.4 | 0.4 M NaCitrate, 150 mMNaCl |
| 6.0, 6.4 | 0.4 M NaCacodylate, 150 mMNaCl |
| 7.0 | 0.4 M NaPhospate, 150 mMNaCl |
| 7.4 | 0.4 M NaHEPES, 150 mMNaCl |
| 8.0, 8.4 | 0.4 M Tris/HCl, 150 mMNaCl |
| 9.0, 9.4 | 0.4 M NaBorate, 150 mMNaCl |

**Supplemental Table 2:** Buffers used during analytical gel filtration experiments.

| pH | Buffer |
| --- | --- |
| 3.0 | 20 mMGlycine/HCl, 150 mMNaCl |
| 5.0 | 20 mMNaCitrate, 150 mMNaCl |
| 7.1 | 20 mMNaHEPES, 150 mMNaCl |
| 8.6 | 20 mMTris/HCl, 150 mMNaCl |

**Figure S1:**
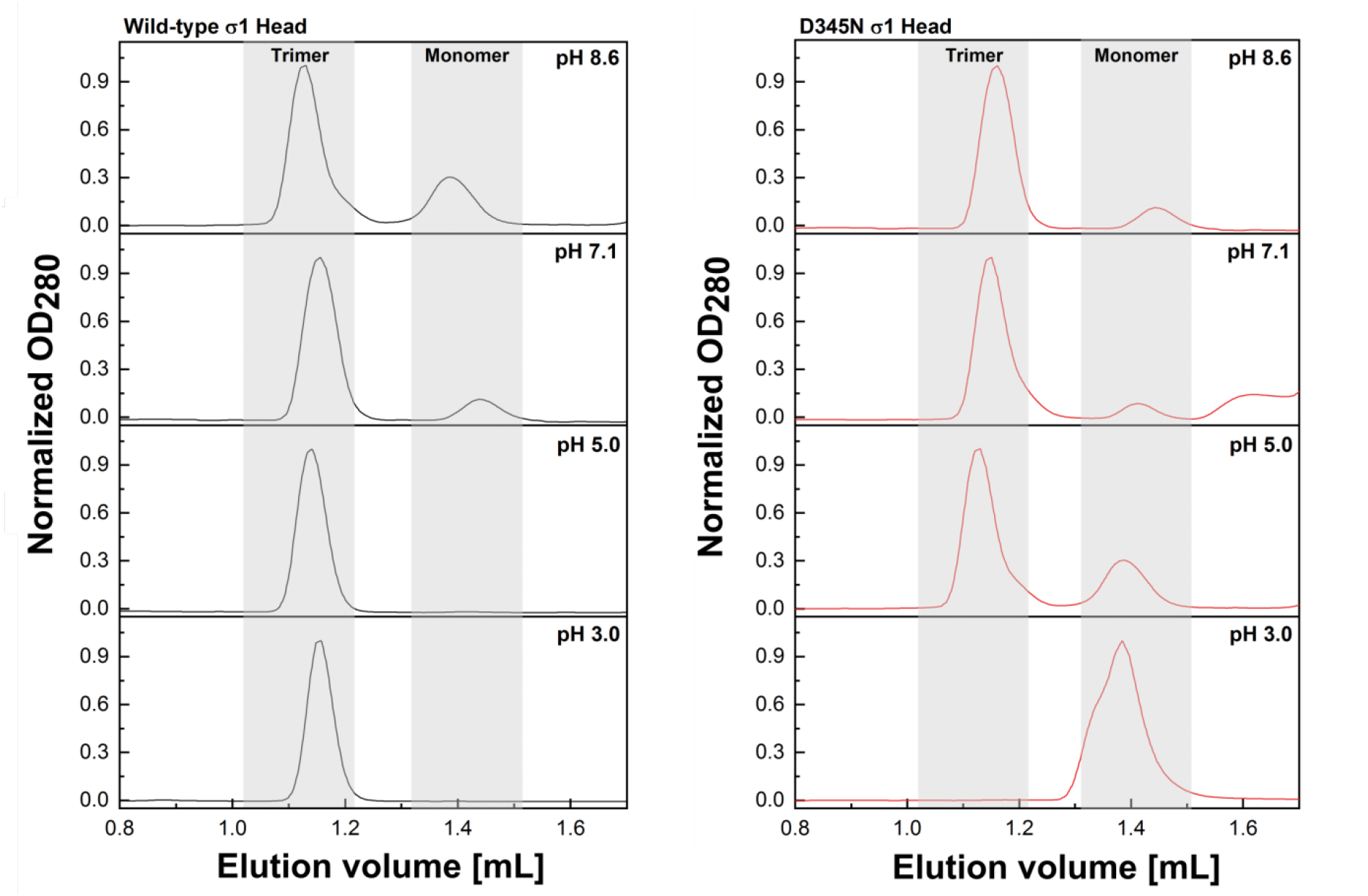
Analytical size exclusion traces of T3D wild-type (black) and D345N (red) σ1 Head samples at pH 3.0, 5.0, 7.1, and 8.6. Absorbance peaks corresponding to trimers and monomers of T3D σ1 Head were integrated in Unicorn and are highlighted by a grey background.

## REFERENCES

1. Selb B, Weber B. 1994. A study of human reovirus IgG and IgA antibodies by ELISA and western blot. J Virol Methods 47:15–25.

2. Tai JH, Williams JV, Edwards KM, Wright PF, Crowe JE, Jr., Dermody TS. 2005. Prevalence of reovirus-specific antibodies in young children in Nashville, Tennessee. J Infect Dis 191:1221–4.

3. Giordano MO, Martinez LC, Isa MB, Ferreyra LJ, Canna F, Pavan JV, Paez M, Notario R, Nates SV. 2002. Twenty year study of the occurrence of reovirus infection in hospitalized children with acute gastroenteritis in Argentina. Pediatr Infect Dis J 21:880–2.

4. Jackson GG, Muldoon RL. 1973. Viruses causing common respiratory infections in man. J Infect Dis 127:328–55.

5. Berger AK, Mainou BA. 2018. Interactions between Enteric Bacteria and Eukaryotic Viruses Impact the Outcome of Infection. Viruses 10.

6. Day JM. 2009. The diversity of the orthoreoviruses: molecular taxonomy and phylogentic divides. Infect Genet Evol 9:390–400.

7. Tyler KL, McPhee DA, Fields BN. 1986. Distinct pathways of viral spread in the host determined by reovirus S1 gene segment. Science 233:770–4.

8. Weiner HL, Powers ML, Fields BN. 1980. Absolute linkage of virulence and central nervous system cell tropism of reoviruses to viral hemagglutinin. J Infect Dis 141:609–16.

9. Weiner HL, Drayna D, Averill DR, Jr., Fields BN. 1977. Molecular basis of reovirus virulence: role of the S1 gene. Proc Natl Acad Sci U S A 74:5744–8.

10. Schelling P, Guglielmi KM, Kirchner E, Paetzold B, Dermody TS, Stehle T. 2007. The reovirus sigma1 aspartic acid sandwich: a trimerization motif poised for conformational change. J Biol Chem 282:11582–9.

11. Kirchner E, Guglielmi KM, Strauss HM, Dermody TS, Stehle T. 2008. Structure of reovirus sigma1 in complex with its receptor junctional adhesion molecule-A. PLoS Pathog 4:e1000235.

12. Barton ES, Forrest JC, Connolly JL, Chappell JD, Liu Y, Schnell FJ, Nusrat A, Parkos CA, Dermody TS. 2001. Junction adhesion molecule is a receptor for reovirus. Cell 104:441–51.

13. Chappell JD, Duong JL, Wright BW, Dermody TS. 2000. Identification of carbohydrate-binding domains in the attachment proteins of type 1 and type 3 reoviruses. J Virol 74:8472–9.

14. Konopka-Anstadt JL, Mainou BA, Sutherland DM, Sekine Y, Strittmatter SM, Dermody TS. 2014. The Nogo receptor NgR1 mediates infection by mammalian reovirus. Cell Host Microbe 15:681–91.

15. Chandran K, Nibert ML. 1998. Protease cleavage of reovirus capsid protein mu1/mu1C is blocked by alkyl sulfate detergents, yielding a new type of infectious subvirion particle. J Virol 72:467–75.

16. Madren JA, Sarkar P, Danthi P. 2012. Cell entry-associated conformational changes in reovirus particles are controlled by host protease activity. J Virol 86:3466–73.

17. Zhang L, Chandran K, Nibert ML, Harrison SC. 2006. Reovirus mu1 structural rearrangements that mediate membrane penetration. J Virol 80:12367–76.

18. Nibert ML, Odegard AL, Agosto MA, Chandran K, Schiff LA. 2005. Putative autocleavage of reovirus mu1 protein in concert with outer-capsid disassembly and activation for membrane permeabilization. J Mol Biol 345:461–74.

19. Fraser RD, Furlong DB, Trus BL, Nibert ML, Fields BN, Steven AC. 1990. Molecular structure of the cell-attachment protein of reovirus: correlation of computer-processed electron micrographs with sequence-based predictions. J Virol 64:2990–3000.

20. Furlong DB, Nibert ML, Fields BN. 1988. Sigma 1 protein of mammalian reoviruses extends from the surfaces of viral particles. J Virol 62:246–56.

21. Dietrich MH, Ogden KM, Long JM, Ebenhoch R, Thor A, Dermody TS, Stehle T. 2018. Structural and Functional Features of the Reovirus sigma1 Tail. J Virol 92.

22. Reiter DM, Frierson JM, Halvorson EE, Kobayashi T, Dermody TS, Stehle T. 2011. Crystal structure of reovirus attachment protein sigma1 in complex with sialylated oligosaccharides. PLoS Pathog 7:e1002166.

23. Prota AE, Campbell JA, Schelling P, Forrest JC, Watson MJ, Peters TR, Aurrand-Lions M, Imhof BA, Dermody TS, Stehle T. 2003. Crystal structure of human junctional adhesion molecule 1: implications for reovirus binding. Proc Natl Acad Sci U S A 100:5366–71.

24. Paul RW, Choi AH, Lee PW. 1989. The alpha-anomeric form of sialic acid is the minimal receptor determinant recognized by reovirus. Virology 172:382–5.

25. Chappell JD, Gunn VL, Wetzel JD, Baer GS, Dermody TS. 1997. Mutations in type 3 reovirus that determine binding to sialic acid are contained in the fibrous tail domain of viral attachment protein sigma1. J Virol 71:1834–41.

26. Reiss K, Stencel JE, Liu Y, Blaum BS, Reiter DM, Feizi T, Dermody TS, Stehle T. 2012. The GM2 glycan serves as a functional coreceptor for serotype 1 reovirus. PLoS Pathog 8:e1003078.

27. Dryden KA, Wang G, Yeager M, Nibert ML, Coombs KM, Furlong DB, Fields BN, Baker TS. 1993. Early steps in reovirus infection are associated with dramatic changes in supramolecular structure and protein conformation: analysis of virions and subviral particles by cryoelectron microscopy and image reconstruction. J Cell Biol 122:1023–41.

28. Nibert ML, Chappell JD, Dermody TS. 1995. Infectious subvirion particles of reovirus type 3 Dearing exhibit a loss in infectivity and contain a cleaved sigma 1 protein. J Virol 69:5057–67.

29. Dietrich MH, Ogden KM, Katen SP, Reiss K, Sutherland DM, Carnahan RH, Goff M, Cooper T, Dermody TS, Stehle T. 2017. Structural Insights into Reovirus sigma1 Interactions with Two Neutralizing Antibodies. J Virol 91.

30. Koehler M, Aravamudhan P, Guzman-Cardozo C, Dumitru AC, Yang J, Gargiulo S, Soumillion P, Dermody TS, Alsteens D. 2019. Glycan-mediated enhancement of reovirus receptor binding. Nat Commun 10:4460.

31. Chappell JD, Prota AE, Dermody TS, Stehle T. 2002. Crystal structure of reovirus attachment protein sigma1 reveals evolutionary relationship to adenovirus fiber. EMBO J 21:1–11.

32. Campbell JA, Schelling P, Wetzel JD, Johnson EM, Forrest JC, Wilson GA, Aurrand-Lions M, Imhof BA, Stehle T, Dermody TS. 2005. Junctional adhesion molecule a serves as a receptor for prototype and field-isolate strains of mammalian reovirus. J Virol 79:7967–78.

33. Cavalli A, Prota AE, Stehle T, Dermody TS, Recanatini M, Folkers G, Scapozza L. 2004. A molecular dynamics study of reovirus attachment protein sigma1 reveals conformational changes in sigma1 structure. Biophys J 86:3423–31.

34. Morin MJ, Warner A, Fields BN. 1994. A pathway for entry of reoviruses into the host through M cells of the respiratory tract. J Exp Med 180:1523–7.

35. Wolf JL, Rubin DH, Finberg R, Kauffman RS, Sharpe AH, Trier JS, Fields BN. 1981. Intestinal M cells: a pathway for entry of reovirus into the host. Science 212:471–2.

36. Pye G, Evans DF, Ledingham S, Hardcastle JD. 1990. Gastrointestinal intraluminal pH in normal subjects and those with colorectal adenoma or carcinoma. Gut 31:1355–7.

37. Johansson ME, Larsson JM, Hansson GC. 2011. The two mucus layers of colon are organized by the MUC2 mucin, whereas the outer layer is a legislator of host-microbial interactions. Proc Natl Acad Sci U S A 108 Suppl 1:4659–65.

38. He G, Shankar RA, Chzhan M, Samouilov A, Kuppusamy P, Zweier JL. 1999. Noninvasive measurement of anatomic structure and intraluminal oxygenation in the gastrointestinal tract of living mice with spatial and spectral EPR imaging. Proc Natl Acad Sci U S A 96:4586–91.

39. Nagita A, Amemoto K, Yoden A, Aoki S, Sakaguchi M, Ashida K, Mino M. 1996. Diurnal variation in intragastric pH in children with and without peptic ulcers. Pediatr Res 40:528–32.

40. Parks TD, Leuther KK, Howard ED, Johnston SA, Dougherty WG. 1994. Release of proteins and peptides from fusion proteins using a recombinant plant virus proteinase. Anal Biochem 216:413–7.

41. Oshaben KM, Salari R, McCaslin DR, Chong LT, Horne WS. 2012. The native GCN4 leucine-zipper domain does not uniquely specify a dimeric oligomerization state. Biochemistry 51:9581–91.

42. O’Shea EK, Klemm JD, Kim PS, Alber T. 1991. X-ray structure of the GCN4 leucine zipper, a two-stranded, parallel coiled coil. Science 254:539–44.

43. Studier FW. 2014. Stable expression clones and auto-induction for protein production in E. coli. Methods Mol Biol 1091:17–32.

44. Niesen FH, Berglund H, Vedadi M. 2007. The use of differential scanning fluorimetry to detect ligand interactions that promote protein stability. Nat Protoc 2:2212–21.

45. Karshikoff A, Nilsson L, Ladenstein R. 2015. Rigidity versus flexibility: the dilemma of understanding protein thermal stability. FEBS J 282:3899–917.

46. Herschlag D, Pinney MM. 2018. Hydrogen Bonds: Simple after All? Biochemistry 57:3338–3352.

47. Gerlt JA, Kreevoy MM, Cleland W, Frey PA. 1997. Understanding enzymic catalysis: the importance of short, strong hydrogen bonds. Chem Biol 4:259–67.

48. Frey PA. 1995. Low-barrier hydrogen bonds. Science 268:189.

49. Cleland WW, Frey PA, Gerlt JA. 1998. The low barrier hydrogen bond in enzymatic catalysis. J Biol Chem 273:25529–32.

50. Lin J, Pozharski E, Wilson MA. 2017. Short Carboxylic Acid-Carboxylate Hydrogen Bonds Can Have Fully Localized Protons. Biochemistry 56:391–402.

51. Seiradake E, Cusack S. 2005. Crystal structure of enteric adenovirus serotype 41 short fiber head. J Virol 79:14088–94.

52. Seiradake E, Lortat-Jacob H, Billet O, Kremer EJ, Cusack S. 2006. Structural and mutational analysis of human Ad37 and canine adenovirus 2 fiber heads in complex with the D1 domain of coxsackie and adenovirus receptor. J Biol Chem 281:33704–16.

53. Rafie KL, A.; Fuchs, J.; Rajan, A.; Arnberg, N.; Carlson, LA. 2021. The structure ofenteric human adenovirus 41—A leading cause ofdiarrhea inchildren. Science Advances

54. Hibbert FE, J. 1990. Hydrogen Bonding and Chemical Reactivity. Advances in Physical Organic Chemistry 26:255–379.

55. Stettner E, Dietrich MH, Reiss K, Dermody TS, Stehle T. 2015. Structure of Serotype 1 Reovirus Attachment Protein sigma1 in Complex with Junctional Adhesion Molecule A Reveals a Conserved Serotype-Independent Binding Epitope. J Virol 89:6136–40.

56. Diller JR, Halloran SR, Koehler M, Dos Santos Natividade R, Alsteens D, Stehle T, Dermody TS, Ogden KM. 2020. Reovirus sigma1 Conformational Flexibility Modulates the Efficiency of Host Cell Attachment. J Virol 94.

57. Bangaru S, Lang S, Schotsaert M, Vanderven HA, Zhu X, Kose N, Bombardi R, Finn JA, Kent SJ, Gilchuk P, Gilchuk I, Turner HL, Garcia-Sastre A, Li S, Ward AB, Wilson IA, Crowe JE, Jr. 2019. A Site of Vulnerability on the Influenza Virus Hemagglutinin Head Domain Trimer Interface. Cell 177:1136–1152 e18.

58. Zost SJ, Gilchuk P, Case JB, Binshtein E, Chen RE, Reidy JX, Trivette A, Nargi RS, Sutton RE, Suryadevara N, Williamson LE, Chen EC, Jones T, Day S, Myers L, Hassan AO, Kafai NM, Winkler ES, Fox JM, Steinhardt JJ, Ren K, Loo YM, Kallewaard NL, Martinez DR, Schafer A, Gralinski LE, Baric RS, Thackray LB, Diamond MS, Carnahan RH, Crowe JE. 2020. Potently neutralizing human antibodies that block SARS-CoV-2 receptor binding and protect animals. bioRxiv doi:10.1101/2020.05.22.111005.

59. Zost SJ, Gilchuk P, Chen RE, Case JB, Reidy JX, Trivette A, Nargi RS, Sutton RE, ! Suryadevara N, Chen EC, Binshtein E, Shrihari S, Ostrowski M, Chu HY, Didier JE, I MacRenaris KW, Jones T, Day S, Myers L, Lee FE, Nguyen DC, Sanz I, Martinez I DR, Baric RS, Thackray LB, Diamond MS, Carnahan RH, Crowe JE. 2020. Rapid isolation and profiling of a diverse panel of human monoclonal antibodies targeting the SARS-CoV-2 spike protein. bioRxiv doi:10.1101/2020.05.12.091462.

60. Zhou T, Tsybovsky Y, Gorman J, Rapp M, Cerutti G, Chuang GY, Katsamba PS, Sampson JM, Schon A, Bimela J, Boyington JC, Nazzari A, Olia AS, Shi W, Sastry M, Stephens T, Stuckey J, Teng IT, Wang P, Wang S, Zhang B, Friesner RA, Ho DD, Mascola JR, Shapiro L, Kwong PD. 2020. Cryo-EM Structures of SARS-CoV-2 Spike without and with ACE2 Reveal a pH-Dependent Switch to Mediate Endosomal ! Positioning of Receptor-Binding Domains. Cell Host Microbe 28:867–879 e5.

